# Phylogenomics and effector analysis of *Plasmodiophora brassicae* genomes unveils a unique and highly divergent clade in Australia

**DOI:** 10.1101/2025.07.30.667784

**Authors:** Maxim Prokchorchik, Matthew A Campbell, Peri Tobias, Jacob Downs, Sera Choi, Brent N. Kaiser

**Affiliations:** Centre for Carbon, Water and Food, School of Life and Environmental Sciences, Brownlow Hill, NSW, University of Sydney, Australia; School of Life and Environmental Sciences, The University of Sydney, Camperdown, NSW, Australia; Plant Breeding Institute, School of Life and Environmental Sciences, Cobbitty, NSW, Australia

## Abstract

Clubroot, caused by the protist pathogen *Plasmodiophora brassicae*, is a soilborne disease that leads to significant yield losses in a broad range of brassica crops worldwide. Despite its significant impact on agriculture, genomic and molecular studies of clubroot disease are hindered due to the complex life cycle and the obligate biotrophic nature of the pathogen, which prevents its cultivation *in vitro*. In addition, several genotypes of *P. brassicae* are often present as a mixture in a single field sample, making it challenging to resolve high-quality assemblies of individual genomes. Recently, several genomes were assembled for some strains of *P. brassicae*, however, in-depth genomic analysis of this devastating pathogen remains limited.

In this study, we generated complete telomere-to-telomere genome assemblies for three Australian *P. brassicae* field isolates using PacBio Revio HiFi sequencing combined with chromatin conformation capture technology (Hi-C). Importantly, we could generate two nearly fully phased haplotype genome assemblies from one of the field isolates. Additionally, we sequenced 14 isolates of *P. brassicae,* sampled from a wide geographical range across Australia using Illumina technology and performed an in-depth global comparative phylogenetic analysis. We revealed that Australian isolates are classified into three different clades, including a highly divergent and unique clade present only in Australia. Comparative analysis showed that the predicted effector profile of the Australian-unique clade is distinct from other clades, further supporting phylogenetic divergence. Altogether, we demonstrate the first haplotype-resolved genome of Australian *P. brassicae* isolates from a single clubroot field sample, and provide an in-depth phylogenetic analysis of global and Australian isolates. These findings, together with core effector profile analysis, will further advance the research to combat this destructive pathogen.

## Introduction

Clubroot of brassica crops, caused by an obligate biotrophic protist *Plasmodiophora brassicae* (Woronin 1877), is a devastating plant disease leading to significant crop yield loss worldwide. *P. brassicae* is a soilborne plant pathogen that enters the plant roots through root hairs and leads to deformation of the entire root, forming tumour-like clubs during disease development. These clubs, produced mainly by uncontrollable cell division, act as a nutrient sink and eventually lead to stunted plant growth and plant death in some cases [1]. *P. brassicae* was initially classified as a member of the fungal *Myxomycetes* group, then later reassigned to the eukaryotic *Rhizaria* group based on its distinctive physiology and DNA sequence similarity [2–4].

The host range of *P. brassicae* presumably covers nearly all *Brassica* species, including many cultivated crops and wild species [5, 6]. In addition to its wide host range, the dissemination of this disease between fields is rapid, leading to disease resistance breakdown in previously resistant crop varieties [7]. Recently, a severe outbreak of clubroot disease in a previously clubroot-resistant commercial canola cultivar has been reported in Canada, leading to significant damage to the Canadian canola industry [8]. Once the field is infected, it is difficult to eradicate the pathogen since the resting spores of *P. brassicae* can retain viability in soil for up to 20 years [9]. Moreover, the geographical distribution of *P. brassicae* has been constantly expanding, probably associated with human migration and agricultural activities, resulting in clubroot infections being reported in more than 80 countries across Europe, America, Asia and Oceania [1]. Despite its devastating impact on a wide range of brassica crops worldwide, genomic studies on *P. brassicae* remain limited and have only recently started to emerge.

One of the great challenges in *P. brassicae* genomic studies is its complex obligate biotrophic life cycle, making isolation of a single pathotype from infected plant roots problematic. *P. brassicae* often exist as a mixture of genotypes, even within a single club, therefore a tedious procedure of single spore isolation is required before genomic DNA extraction for sequencing [10]. The latest phylogenomic analysis of *P. brassicae* isolates indicated that there are five clades that did not correlate with experimentally determined pathotypes [11]. Recently, the first telomere-to-telomere (T2T) genome of Canadian *P. brassicae* isolate Pb3A from canola was reported [12], laying a foundation for large comparative genomic studies. Higher-quality genomes from a larger number of isolates originating from different geographic regions will help our understanding of the evolution and potential migration trajectories of clubroot. While there is genome information for *P. brassicae* isolates from countries in Europe, America and Asia, there are none available for countries in Oceania, such as Australia, where canola cultivation has recently expanded as an important export commodity [10, 13–16].

Pathogens secrete small proteins called effectors into host plants to suppress their immune system and manipulate the host physiology to proliferate and cause disease [17]. Resistant plants, in turn, can recognise these effectors by resistance (R) proteins and establish effector-triggered immunity (ETI) [17]. *P. brassicae* is reported to encode approximately 600 effectors [18, 19]. Function in virulence has been characterised for only a few effectors, while no effector-*R* gene pair has been reported [18, 20–24]. Recent improvement of protein structure modelling tools allowed prediction of structurally conserved effectors likely playing a similar role in virulence [25], however, large-scale systematic comparative analysis of effectors from different *P. brassicae* isolates is absent.

Here, we report the first complete T2T and nearly fully haplotype-resolved assemblies of Australian field-isolated *P. brassicae* using HiFi Revio long-read and Hi-C technologies. We sampled and sequenced 14 additional isolates, using Illumina technology, from across Australia and incorporated this data into a global phylogeny of *P. brassicae* isolates. Strikingly, we observed an Australian-specific clade highly divergent from other isolates. In addition, we conducted comparative effector profile analysis in different sets of *P. brassicae* isolates and identified core effectors shared between different isolates. Finally, we systematically compared the effector profiles of representative isolates from each clade, defined from the phylogenomic analysis, and discovered that *P. brassicae* isolates in the divergent Australian-specific clade possess a distinct set of effectors. Altogether, our research advances the understanding of *P. brassicae* evolutionary patterns and provides a foundation resource for further research of this pathogen’s virulence strategies.

## Results and Discussion

### Sequencing the haplotypes from clubroot isolates using chromatin capture (Hi-C)

As *P. brassicae* cannot be cultured outside the appropriate plant host, extracting pure and high molecular weight genomic DNA for long-read sequencing requires large amounts of infected plant material. To identify the susceptible plant host for Australian *P. brassicae* isolate propagation to obtain a sufficient amount of genomic DNA, we infected several brassica crop species, including mizuna (*Brassica rapa* var. *niposinica*), broccoli (*B. oleracea* var. *italica*), Ethiopian mustard (*B. carinata*), Chinese cabbage (*B. rapa* subsp. *pekinensis*) and kale (*B. oleracea* var. *viridis*) with resting spores extracted from infected brassica plants collected from Lindenow (here referred to as VIC3, Supplementary Table 1). While VIC3 successfully infected all tested species, confirming its virulence in a wide range of *Brassica* species, similarly to other reported *P. brassicae* isolates [26, 27], mizuna showed the most severe symptoms (Supplementary Figure S1A). We have further confirmed that the other two Australian field isolates, VIC1 and VIC2, collected in Maffra (Supplementary Table 1), cause strong symptoms in mizuna plants as well (Supplementary Figure S1B). Therefore, we chose mizuna to use as a host throughout the experiments in this study.

To increase the fraction of *P. brassicae* genomic DNA extracted from the infected plants, we first purified and concentrated the resting spores from the infected roots using Ficoll gradient centrifugation [28]. A total of three field isolates (VIC1, VIC2 and VIC3) were sequenced using PacBio HiFi long-read sequencing technology, while VIC2 was also sequenced with chromatin conformation capture (Hi-C) technology. Using these datasets, we assembled 20 complete T2T (telomere-to-telomere) chromosomes for each of the isolates and a nearly complete second haplotype for VIC2 (Table 1). We have designated the primary VIC2 assembly as VIC2h1, consisting of 20 T2T chromosomes and having a BUSCO score of 88.24%, while the secondary phased genome was designated as VIC2h2, with 16 T2T chromosomes, 4 nearly complete chromosomes missing one of the telomeres, and a BUSCO score of 85.88% (Table 1). We have further confirmed that the BUSCO score remains a poor metric for assessing plant pathogenic protists genome assemblies, achieving a similar level of completeness to the genome reported recently [12] (Fig 1A). To improve gene annotation prediction, we further performed RNA-seq transcriptomic analysis for the samples extracted from purified resting spores and infected plants (21 days post-inoculation) to achieve high sequencing coverage, especially for genes differentially expressed during different life stages of the pathogen. As a result, we increased the number of predicted protein-coding genes to 11,103 (Table 1), in comparison to the latest Pb3A annotation [12].

**Fig 1.**
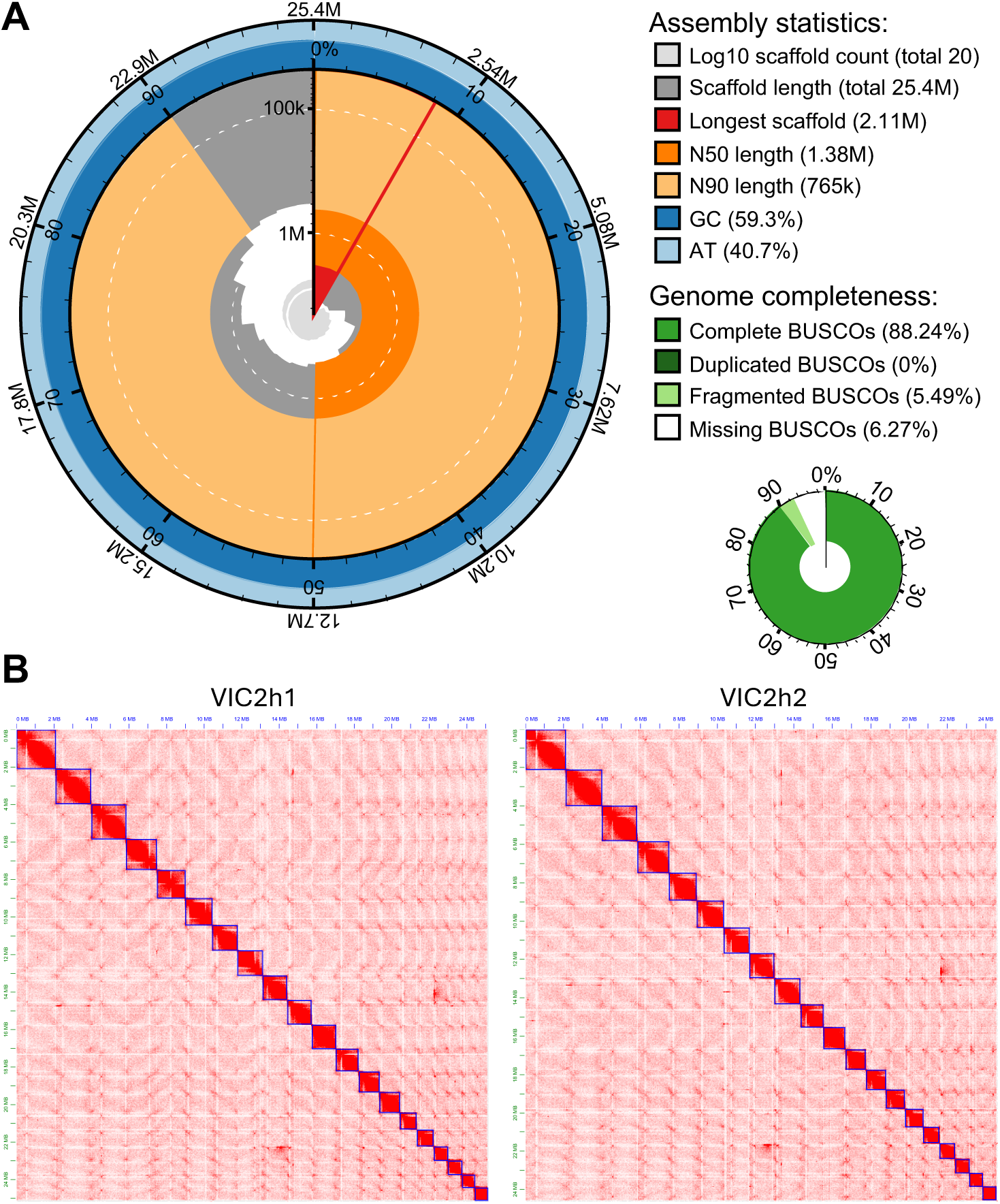
Assembly statistics for the Australian Plasmodiophora brassicae reference genome. (A) Snail plot showing the key statistics for the VIC2h1 complete T2T genome assembly. (B) Contact maps showing the Hi-C support for the VIC2h1 and VIC2h2 haplotype scaffolding.

**Table 1.** Assembly statistics of the complete *P. brassicae* genome.

|  | **Pb3A [18]** | **VIC1** | **VIC2h1** | **VIC2h2** | **VIC3** |
| --- | --- | --- | --- | --- | --- |
| Genome size (bp) | 25,283,981 | 25,498,282 | 25,409,529 | 24,706,353 | 25,446,875 |
| Number of scaffolds / chromosomes (T2T) | 20/20 | 20/20 | 20/20 | 20/16 | 20/20 |
| Largest scaffold (bp) | 2,102,640 | 2,110,470 | 2,110,616 | 2,147,489 | 2,110,644 |
| N50 length (bp) | 1,371,253 | 1,383,291 | 1,383,171 | 1,337,268 | 1,382,327 |
| N90 length (bp) | 747,440 | 765,188 | 765,189 | 770,335 | 765,309 |
| GC content (%) | 59.49 | 59.31 | 59.3 | 59.28 | 59.31 |
| Genes predicted | 10,521 | 11,334 | 11,103 | 10,801 | 11,331 |
| Repeat content (%) | 11.88 | 15.70 | 15.49 | 15.71 | 15.55 |
| Complete BUSCOs (%) | 88.2 | 88.24 | 88.24 | 85.88 | 88.24 |
| Duplicated BUSCOs (%) | 0 | 0 | 0 | 0 | 0 |
| Fragmented BUSCOs (%) | 5.1 | 5.49 | 5.49 | 5.1 | 5.49 |
| Missing BUSCOs (%) | 6.7 | 6.27 | 6.27 | 9.02 | 6.27 |

In many cases, the field collections of *P. brassicae* constitute a mixture of different genotypes. By using the properties of Hi-C data with the highly accurate HiFi reads, at least two diverse, nearly complete genome assemblies from the same sample were resolved (Fig 1B). These results suggest that the improvement of HiFi read accuracy, combined with the increasing affordability of Hi-C data present a feasible method to capture the diversity of *P. brassicae* haplotypes from field isolates, enabling quick phylogenetic and composition profiling of the mixed field collections.

### Whole genome comparison reveals large genomic rearrangements between P. brassicae strains

We next sought to compare the complete T2T and haplotype-phased Australian *P. brassicae* genome, here named VIC2h1, to the recently published Canadian isolate Pb3A [12]. Interestingly, we identified several large genomic rearrangements, including 14 inversions (spanning over 136,877 bp) and 119 translocations (spanning over 361,071 bp) (Fig 2A). We further discovered 428 duplications in the VIC2h1 genome spanning over 895,856 bp and 318 duplications in the Pb3A genome spanning over 644,051 bp.

**Fig 2.**
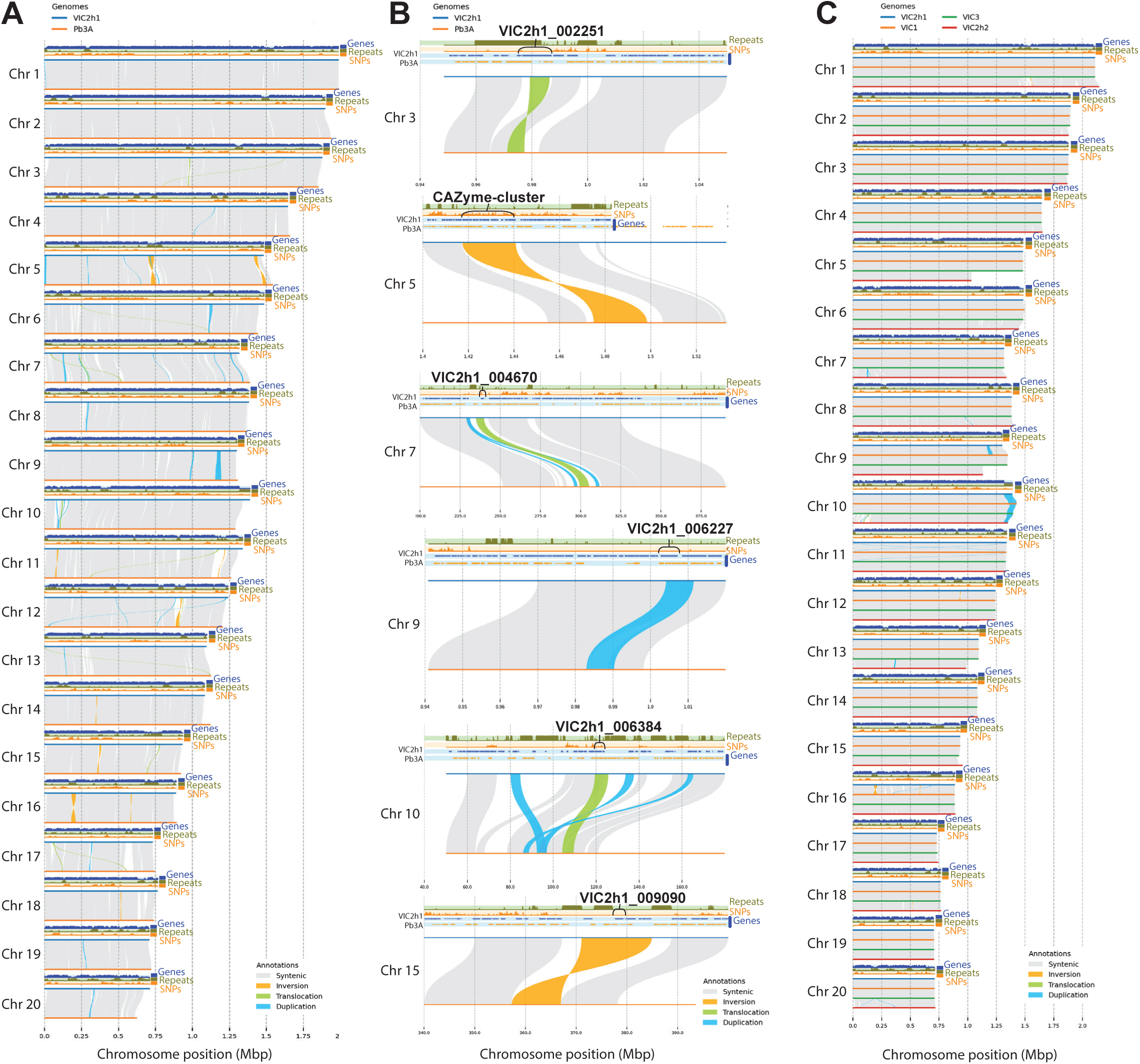
Whole-genome comparison shows the large genomic rearrangements in the complete P. brassicae genomes. (A) Whole-genome comparison between Australian VIC2h1 and Canadian Pb3A genomes. (B) Large structural rearrangements affecting the CAZyme-cluster and putative secreted protein coding genes. (C) Whole-genome comparison between complete Australian genomes. All genome comparisons were carried out using MUMmer and visualised using SyRi and plotsr. Only rearrangements larger than 2kb were plotted. Genes, repeats, and SNP tracks were visualised with a 10,000 nt sliding window.

Pathogen genome rearrangements are often related to their pathogenicity [29, 30]. Therefore, we next investigated if these genome rearrangements are associated with genes potentially important for the pathogen’s virulence (effectors and carbohydrate-active enzymes (CAZymes) [31]. Importantly, we observed that out of the 22 largest rearrangements, at least five were affecting the putative effector genes and one large CAZyme cluster (Fig 2B). Furthermore, predicted effectors, including VIC2h1_002251, VIC2h1_004670, VIC2h1_006227, VIC2h1_006384, VIC2h1_009090, that are affected in the rearrangements seem to be associated with a variety of transposable elements, such as DNA/TcMar-Tc2 repeat, LTR, and DNA/PiggyBac-X (Fig 2B).

To assess the diversity of the *P. brassicae* genomes derived from a single collection, we have performed the genomic comparison between VIC2h1 and VIC2h2 haplotype-resolved genomes. As expected, these genomes showed significantly less variation in comparison to the Pb3A genotype. We could identify 8 inversions (spanning over 52,384 bp), 18 translocations (spanning over 44,506 bp), and 176 duplications (spanning over 339,936 bp) (Fig 2C). These results suggest that the local complete genomes of currently circulating isolates found in Australia are more closely related to each other, than to Pb3A.

To check the similarity of the complete Australian assemblies, we have performed whole-genome comparisons between VIC1, VIC2h1, VIC2h2 and VIC3 assemblies. Interestingly, the VIC2h2 genome showed the farthest relation to all the other assemblies (Fig 2C). This highlights that Hi-C support was important to phase the genomes from the same sample, while in its absence, the diversity of genomes may have collapsed during the assembly process.

### Phylogenetic analysis of P. brassicae isolates reveals a unique clade found only in Australia

While there is sequence information for *P. brassicae* isolates from different countries, until now there was no data for any of the Australian isolates. To understand the local diversity of the *P. brassicae* isolates in Australia, we sequenced a representative set of collections from a wide geographic range across Australia. Using the specimens from the New South Wales Department of Primary Industries (NSW DPI) herbarium and field collections from the brassica vegetable farms, we managed to generate a set of 17 samples covering the South Australia (SA), New South Wales (NSW), Tasmania (TAS) and Victoria (VIC) states of Australia (Supplementary Table 1) (Fig 3A). Among them, we successfully sequenced 14 herbarium specimens using NGS Illumina technology and generated draft genome assemblies (Supplementary Table 2).

**Fig 3.**
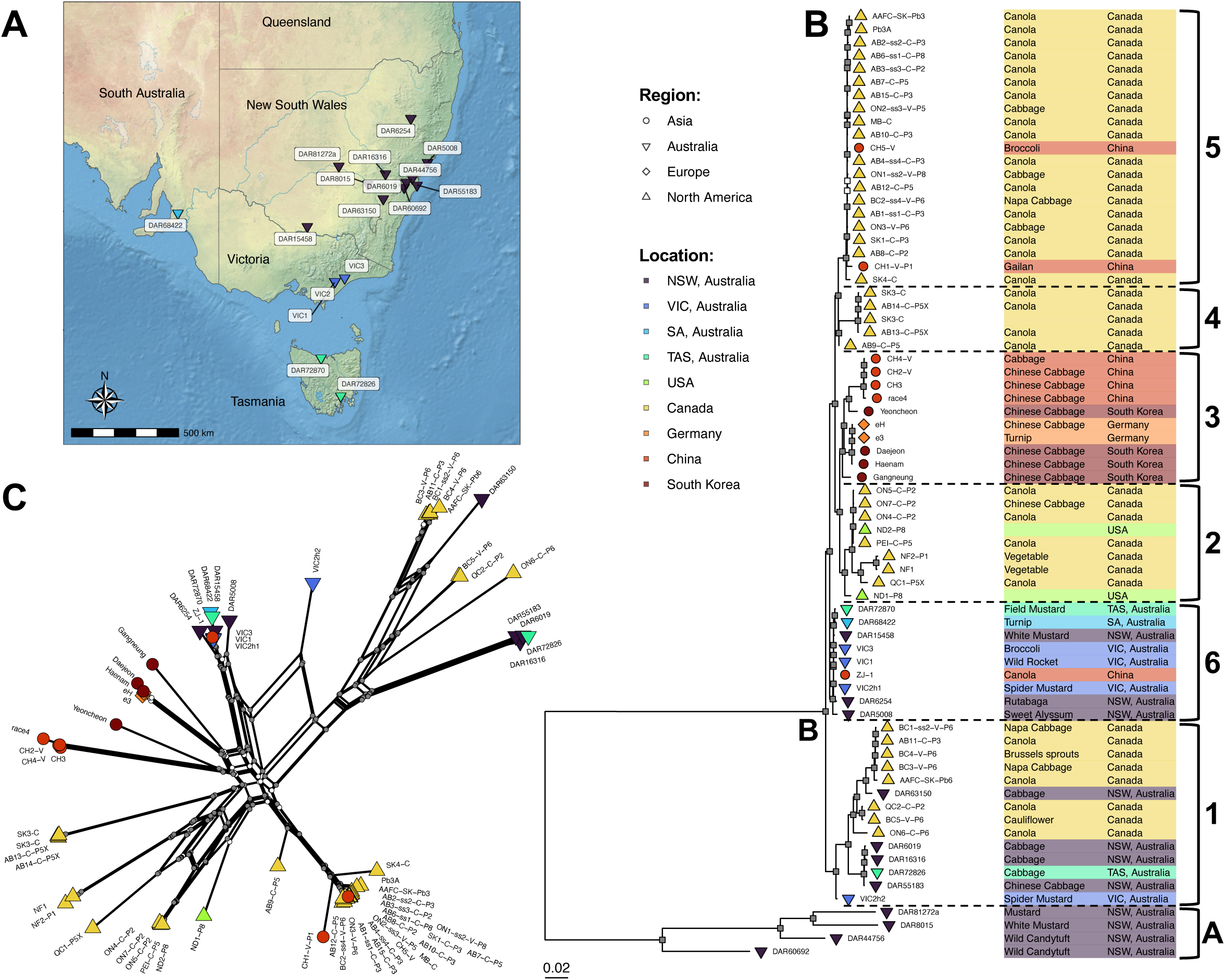
Phylogenetic analysis of P. brassica isolates. (A) Geographic distribution of the sequenced *P. brassicae* isolates across Australia. (B) Phylogenetic tree and (C) network analysis of the available and newly sequenced *P. brassicae* isolates. Core genome alignment for (A) and (B) was performed with the PhaME pipeline, and a tree was built using IQ-Tree2. Only branches with bootstrap support of more than 70% are shown. Nodes with bootstrap support values > 75% but <= 90% are white and nodes with bootstrap support values > 90% are grey in both panels (B) and (C).

Previously, the *P. brassicae* isolates were reported to form 5 phylogenetic clades [11]. We assessed the evolutionary relationship of 72 global *P. brassicae* isolates, consisting of 51 assemblies available from the NCBI database, 17 Australian (sequenced in this study) and four South Korean assemblies generated from publicly available short-read data [32]. The multiple sequence alignment (MSA) used to build the phylogenetic tree was 109,960 sites in length, with 81,926 parsimony-informative sites and 13,228 distinct alignment patterns. A best fit model of TVM+F+ASC+R4 was chosen by BIC and used for the ML tree search. The ML phylogeny indicates two major clades separated by a long branch (Fig 3B). A clade of DAR81272a, DAR8015, DAR44756 and DAR60692 showed high divergence from any *P. brassicae* sequenced to date and highly supported monophyly (BS=100%) (Fig 3B). This clade, geographically restricted to the central NSW region, was found in white mustard (*Sinapis alba*) and candytuft plants (*Iberis amara*) and is referred to as clade A going forward (Fig 3B). The remainder of the Australian assemblies were placed into a second, maximally supported clade – clade B – with collections of both wide geographic and host range (Fig 3B). All the complete T2T assemblies generated in this study, except for VIC2h2, form the closely related clade of isolates together with DAR72870, DAR68422, DAR15458, DAR6254 and DAR5008 herbarium specimens. These isolates have a wide geographic range inside Australia and are closely related to the ZJ-1 isolate from China, and we suggest designating this clade as 6 (Fig 3B). The DAR63150, DAR6019, DAR16316, and DAR72826 represent a different clade placed together with multiple Canadian isolates from different hosts that were previously assigned as clade 1 (Fig 3B) [11]. Overall, our phylogenetic analysis agrees with the previously published results [11]. With a larger set of phylogenetically diverse isolates, we have managed to resolve the clades 1 and 2 more precisely and suggest reassigning the QC1-C-P5X, NF1 and NF2-P1 isolates from clade 1 to clade 2.

The phylogenetic network analysis was restricted to clade B (Fig 3C) as the distance between clades A and B is so great that visualisation is problematic. A phylogenetic network analysis of clades A and B is presented in Supplementary Figure S2. The clade B Australian isolates are placed with Canadian or a single Chinese isolate in two groupings (Fig 3C). The VIC2h2 assembly is placed with reticulations between these two larger Australian groupings, indicating that a mixture of genotypes may be present in this sequence (Fig 3C).

Our data supports and reinforces the current understanding of the *P. brassicae* phylogeny and introduces previously undescribed diversity. The phylogenetic distance between the VIC2 haplotypes further shows that different genotypes may be present in the same clubroot in the field. Clade A, with its deep divergence from all other *P. brassicae* genome assemblies, is of unclear origin. One of the potential sources of clade A could be the United Kingdom, from where no genome assemblies are currently available. If clade A is recovered from diversity present in the United Kingdom through additional sequencing, this would support the movement of clade A to Australia by early settlers. Alternatively, the divergence observed may have different origins, such as a native Australian strain, a hypothesis that would be difficult to validate, as proving the absence of clade A outside Australia is challenging.

### Small secreted protein prediction and analysis in Australian and global P. brassicae isolates

Small secreted effector proteins, delivered from the pathogens, are considered the main molecular tools facilitating the manipulation of plant physiology to cause disease [17]. Due to their importance in virulence, effector genes are under high selective pressure, leading to diversification of effector sets in different isolates [33]. To compare the effector profiles in the complete genomes of Australian isolates and previously reported Pb3A, we first carried out effector prediction in VIC1, VIC2h1, VIC2h2, VIC3 and Pb3A. We have defined effectors as small (<400 AA) proteins with predicted signal peptides and no transmembrane domains [34]. We identified 663 putative effectors in the VIC1 genome, 641 and 620 in VIC2h1 and VIC2h2, respectively, 656 in VIC3 and 627 in Pb3A (Fig 4A and B). A smaller number of predicted effectors in VIC2h2 is possibly due to incomplete assembly.

**Fig 4.**
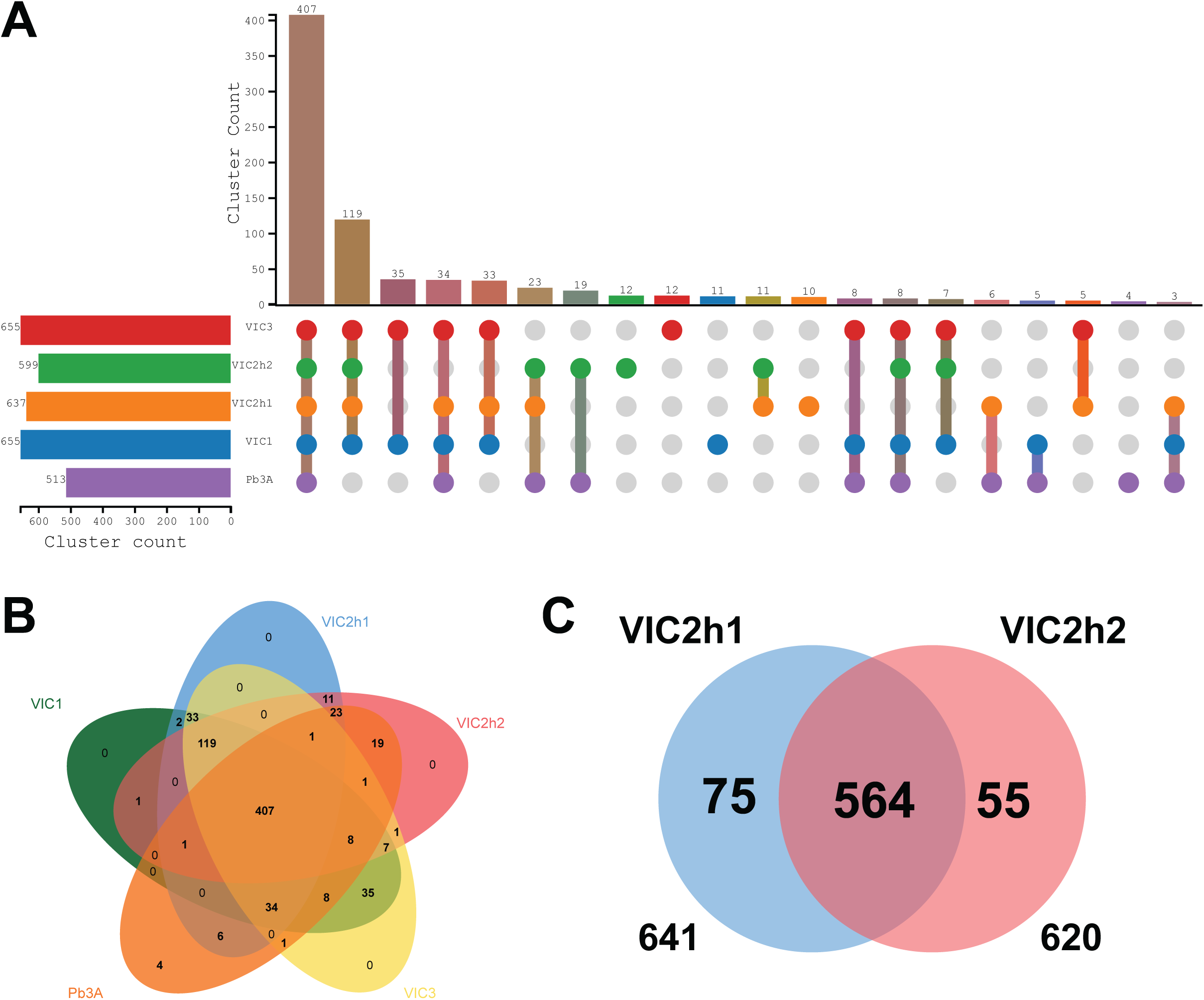
Putative effector analysis in the complete P. brassicae genomes. (A) Upset and (B) Venn diagram plots showing the clusters of putative effectors shared between the VIC1, VIC2h1, VIC2h2, VIC3 and Pb3A complete genomes. (C) Venn diagram showing conserved and unique putative effectors in VIC2h1 and VIC2h2. Effector clusters in all panels were identified using OrthoVenn software with the OrthoMCL algorithm used for cluster discovery.

Despite the diversification pressure on effector genes, the effectors critically important for virulence are often shared between different isolates [35]. To gain comparative insight into effector profiles across the complete genomes, we used orthoMCL to cluster all the effectors from VIC1, VIC2h1, VICh2, VIC3 and Pb3A. A total of 407 orthologous effector clusters shared across all the complete genomes were identified (Fig 4A and B). Interestingly, there were 119 orthologs identified that are unique to the Australian isolates (Fig 4A). Orthologous effectors shared across all the isolates are of particular interest as they are likely to include the conserved core effectors critical for pathogen fitness and/or disease development [35]. In contrast, the 119 orthologous effectors unique to Australian isolates are likely to include the specific ones associated with Australian plant hosts and environmental conditions.

The two haplotype-phased genomes recovered from the VIC2 collection showed considerable phylogenetic distance between each other (Fig 3B). To evaluate how different the putative effector sets they carry are, we clustered the putative effector proteins using orthoMCL. Interestingly, while most putative effectors were conserved between the VIC2h1 and VIC2h2 genomes, a considerable number of 75 putative effector proteins were unique to VIC2h1, while there were 55 putative effectors unique to VIC2h2 (Fig 4C). This data highlights that even in one clubroot sample, there are likely to be several strains of *P. brassicae* expressing different sets of effectors. It would be interesting to investigate if *P. brassicae* can use the pool of effectors secreted from different strains to synergistically infect the host and promote disease development.

The phylogenetic analysis showed that the Australian-specific clade A seems to be genetically distant from all the other clades (Fig 3B). To investigate if effector profiles correlate with and further support the phylogenetic distance between clade A and others, we conducted comparative effector analysis on at least one representative genome from each clade. We predicted and clustered effectors using Orthofinder and generated a heatmap showing the presence/absence of clustered effectors in the representative genomes from each clade (Fig 5). The representative genomes were chosen based on the quality and completeness of the assemblies. Strikingly, we could observe that the presence/absence patterns of the putative effectors follow the phylogenetic delineation closely and there is a clear difference between the effectors sets in clades A and B (Fig 5). We further observed that the putative effector sets in clades 1 and 6 show strong similarity within the respective clades (Fig 5). We have further identified 264 core effector orthologs among these genomes (Fig 5). Together, these findings outline that the difference of the Australian clade A is not only manifested in phylogenetic placement, but also in their diverse sets of putative effectors. These extremely divergent isolates were all virulent on their plant hosts, meaning that the relatively compact core effectorome may be critical to trigger disease. It is noteworthy that the effectors predicted and analysed in this analysis may not represent the complete effector profiles for each strain, due to the different sequencing quality in the datasets used. Meanwhile, the core 407 families effectorome, identified among less divergent, but complete T2T assemblies (Fig 4), can be used as a more reliable dataset for further effector function studies.

**Fig 5.**
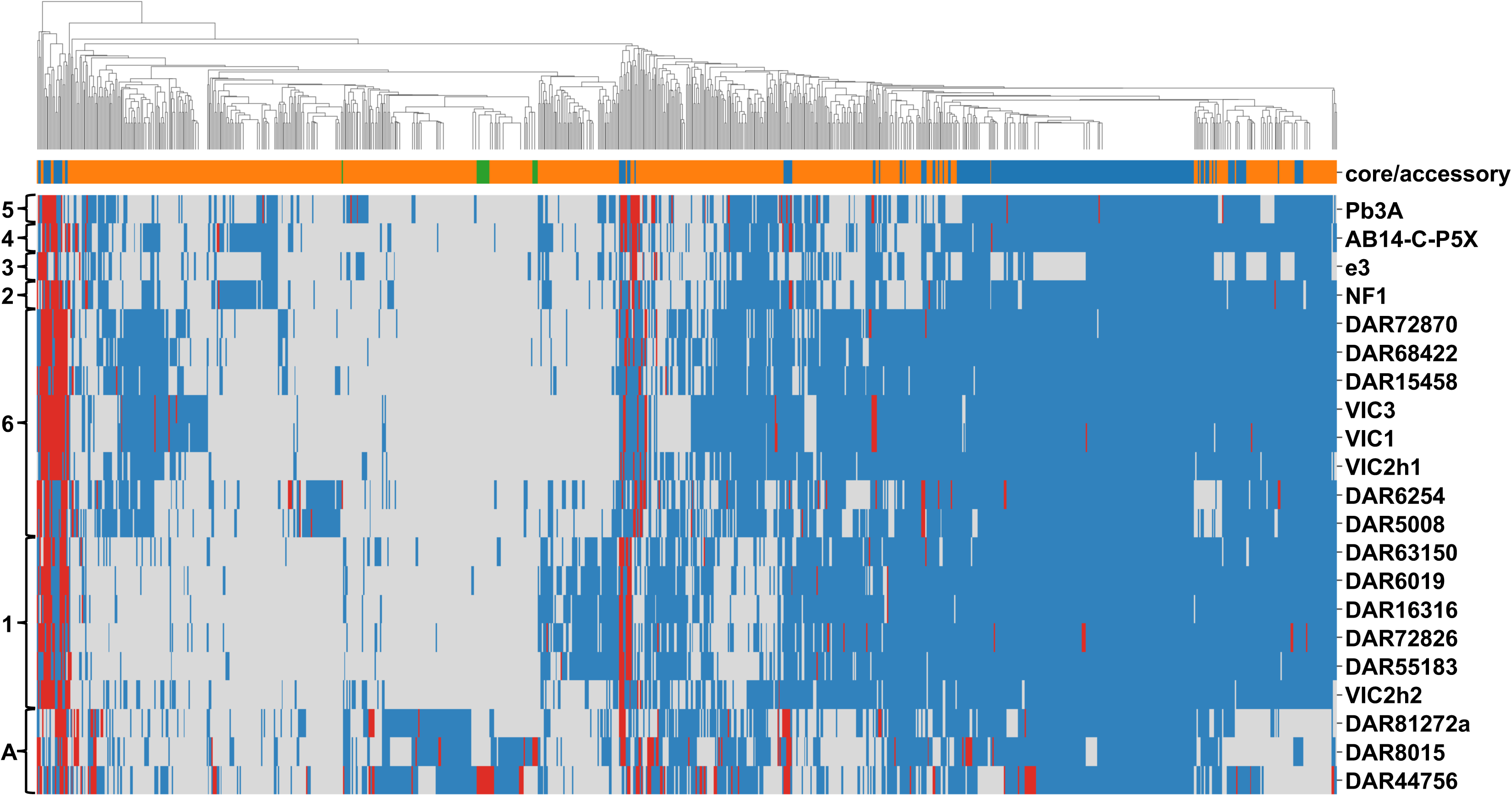
Heatmap showing the orthologous putative effectors shared among the set of Australian isolates and the representatives of each phylogenetic clade. Blue shading in the heatmap plot denotes the presence of a single copy of the effector gene, while red shading denotes the existence of multiple copies. Core effector genes (present in 90% of the genomes) are marked by the blue shading on the core/accessory bar plot, singletons are marked by green and accessory effector genes are shown in orange.

## Materials and methods

### Clubroot isolates handling and plant infection

Viable field collections were sampled from brassica fields in the state of Victoria, Australia (Supplementary Table S1). Herbarium isolates were obtained from DPI (Department of Primary Industries) NSW (New South Wales) plant pathogen herbarium (Supplementary Table S1). Resting spores, representing the haploid uninucleate state for *P. brassicae* [36] were purified from the infected clubroots and reinfected in different brassica host plants. The spore concentration was adjusted to 1 x 10^7^ spores/ml for plant infections. Mizuna (*Brassica rapa* var*. niposinica*) plants were used for routine maintenance of the viable clubroot isolates.

### Clubroot spore purification

*Plasmodiophora brassicae* resting spores were extracted from freshly infected plant roots as described in [10, 37]. In brief, approximately 30 grams of fresh clubbed roots were surface sterilised with Sucrose (10% w/v)/Tween(0.2% v/v)/bleach solution(5% v/v) for 5 min, then washed twice with sterile distilled water. After that, roots were washed with 70% (v/v) ethanol and, finally, with sterile distilled water again. Sterile clubroots were ground using a blender and filtered through 4 layers of Miracloth (Merck, 475855). The resulting mixture was spun down in a Ficoll 400 (GE Healthcare, 17-0300-10) gradient and the interface fraction was washed with distilled water 3 times before further use. The final spore concentration was defined by counting the spores under the microscope using the Neubauer ruled hemocytometer (EMS, 68052-14).

### Nucleic acid extraction

Clean resting spores were centrifuged and concentrated in a minimal volume of MiliQ water. The spore suspension was transferred to the chilled mortar and ground with acid-washed sand in liquid nitrogen. Ground spores were transferred to the 2 mL microcentrifuge tubes and 1 mL of 0.35M D-Sorbitol buffer was added. Spores were washed with 0.35M D-Sorbitol buffer multiple times until the suspension became non-viscous. After that, 800 µL of CTAB extraction buffer with Proteinase K (NEB, P8107S) was added, and samples were incubated at 56°C for 1 h with occasional mixing. After that, RNase A (NEB, T3018L) treatment was performed for 10 min at room temperature, and the mixture was phased using 800 µL of chloroform/isoamyl solution (24/1 (v/v) ratio) (Merck, C0549). The upper phase was transferred to the new tubes, 1 mL of 100% ethanol was added, and DNA was precipitated overnight. After centrifugation, the DNA pellet was washed with 70% (v/v) ethanol and resuspended in 100 µL of TE buffer.

To extract the genomic DNA from the herbarium specimens, dried clubbed roots were soaked in sterile distilled water overnight after surface sterilisation with one wash with Sucrose (10% v/v)/Tween(0.2% v/v)/bleach solution(5% v/v) and one wash with 70% v/v ethanol. Softened tissues were washed thoroughly with 0.2% v/v Tween 20 (Merck, P1379) solution, then rinsed with sterile distilled water and disrupted using a blender. The resulting mixture was filtered through 4 layers of Miracloth (Merck, 475855), but due to the limited amount of tissue, the Ficoll 400 gradient separation was not performed. Instead, spores were thoroughly washed with sterile MiliQ water multiple times and then subjected to DNA extraction.

RNA was extracted from spores or infected roots (at 21 days post infection) using the NEB Monarch RNA extraction kit (NEB, T2110). cDNA was synthesised using the Bio-Rad iScript cDNA Synthesis Kit (Bio-Rad, 1708890), and semi-quantitative PCR with PbACT_F (5’-GGGACATCACCGACTACCTG-3’) and PbACT_R (5’-ACTGCTCCGAGTTGGACAT-3’) primers [38] were used to confirm the presence of the pathogen in the plant samples.

### Sequencing and Hi-C library preparation

HiFi Pac Bio sequencing was performed at the AGRF (Australian Genome Research Facility). DNA libraries for sequencing were prepared using SMRTbell prep kit and sequenced with the SMRT Cell 25M flowcell on the PacBio Revio platform. Crosslinking and DNA extraction for Hi-C analysis were performed using the Phase Genomics Proximo Hi-C (Fungal) Kit according to the manufacturer’s recommendations. Sequencing was performed using the Illumina MiSeq platform at the Biomolecular Research Facility at Australian National University (ANU).

Illumina DNA and RNA sequencing was performed at the AGRF. Genomic DNA libraries were prepared with IDT xGen cfDNA & FFPE Library Preparation Kit and sequenced with Illumina NovaSeq X Plus platform. The libraries for RNA sequencing were prepared with Illumina Stranded mRNA (RNA-Seq) protocol and sequenced with Illumina NovaSeq X Plus platform. To address the potential low quality of DNA extraction from herbarium isolates, all herbaria samples were first sequenced with low coverage, and the clubroot read quality and fraction were estimated using the T2T complete VIC2h1 genome assembly as a reference. Samples with less than 70x coverage were sequenced again to achieve more than 100x coverage among the two runs combined.

### Genome assembly and annotation

To filter out the contaminating non-clubroot reads, raw reads were aligned to the custom database containing all available *Brassicaceae* plant and NCBI bacterial RefSeq genomes using MiniMap2 v2.1 [39] with the ‘-t 24 -ax asm20 --secondary=no’ settings. The non-aligned reads were then aligned to the Pb3A genomic sequence [12] using MiniMap2 to estimate the filtering quality. The sets of all, longer than 20kb, 15kb and 10kb PacBio HiFi reads were assembled using the hifiasm v0.19.8 [40], with ‘-l 3 -hic’ flags where the HiC data was available. The completeness of the resulting draft assemblies was assessed using compleasm v0.2.6 with the eukaryota lineage flag [41]. Each assembly was manually curated to pick the best combination of T2T chromosomes as the whole genome representation. Chromosomes were ordered and oriented according to the recent Pb3A genome assembly [12]. We used RepeatScout v1.0.6 [42] and RepeatModeller v2.0.1 [43] to generate custom species-specific repeat libraries, and then RepeatFinder v4.1.0 [42] was used to annotate repeats in genome assemblies. Soft-masked genomes were used to predict genes using Funannotate v1.8.17 [44] with support of the RNA-seq data aligned to the VIC2h1 reference with Hisat2 v2.1.0 [45].

Illumina reads from the herbarium and four Korean isolates’ genomes were quality filtered and trimmed using FastP v0.23.4 [46]. The resulting reads were assembled using Spades v3.13.0 [47]. Spades-assembled scaffolds were aligned to the VIC2h1 reference genome using MiniMap2 [39] with ‘-t 24 -ax asm20 --secondary=no’ flag. Aligned scaffolds were ordered by length and were assigned as the genome assemblies for the herbarium isolates. Four isolates from Korea (NCBI project: PRJNA476665), for which the raw sequencing data were available at NCBI SRA under SRP150814 accession number, were assembled with the same pipeline.

### Whole-genome comparison and phylogenetic analysis

We used MUMmer v3.1 [48] for comparing whole-genome assemblies of T2T genomes. Large genomic rearrangements were identified with SyRI v1.7.0 [49] and visualised with Plotsr v1.1.1 [50].

*P. brassicae* genome assemblies for phylogenetic analysis were obtained from the NCBI (https://www.ncbi.nlm.nih.gov/datasets/genome/?taxon=37360). Homologous sites for phylogenetic analysis were identified, and a Multiple Sequence Alignment (MSA) was generated with the PhaME pipeline version v1.0.2 [51]. A concatenated Maximum-Likelihood (ML) tree search was conducted with IQ-Tree2 version v2.2.2.6 [52] using the MSA produced by the PhaME pipeline. A best fit model of nucleotide evolution was identified by specifying ModelFinder Plus with ascertainment bias correction and chosen using the Bayesian Information Criterion (BIC) [53]. Support for nodes was evaluated with 10,000 ultrafast bootstrap replicates [54]. The resulting consensus tree file was imported into R and visualised with functions of the ggplot and ggtree packages [55, 56].

As concatenated ML analysis does not model biological realities such as incomplete lineage sorting or hybridisation, we also generated an implicit phylogenetic network to better represent evolutionary processes [57, 58]. Reticulations between branches in this analysis may represent the exchange of genetic material or incomplete lineage sorting but was not an explicated model. The network was created with SplitsTree version v4.19.2 and nodal support assessed with 1,000 bootstraps [58]. Visualisation of the network in R was done with functions of the ggplot, ggtree, phangorn and tanggle packages [55, 59–61].

### Small secreted protein prediction

After genome annotation, the longest transcript for each gene was extracted and translated using the AGAT toolkit using ‘agat_sp_keep_longest_isoform.pl’ command [62]. The resulting proteomes were first analysed for the presence of signal peptides using SignalP6.0 [63]. Then, resulting sequences with signal peptides removed were analysed using deepTMHMM [64], and only the proteins without transmembrane domains were retained. All the putatively secreted proteins were considered in the large genomic rearrangement analysis, while only the ones shorter than 400 AA were considered effectors for clustering and effectorome analysis. For T2T genomes effectorome analysis, we have used the orthoVenn toolkit with the OrthoMCL algorithm for ortholog identification and visualisation [65, 66]. For the large-scale effector presence/absence analysis, we have used Orthofinder v2.5.5[67] and visualised using the seaborn Python library [68].

## Acknowledgments

We would like to acknowledge Dr Jordan Bailey from the New South Wales Department of Primary Industries (NSW DPI) Herbarium for sharing the herbarium specimens. We also would like to acknowledge the contribution of the Plant Pathogen ‘Omics Initiative consortium in the generation of data used in this publication. The Initiative is supported by funding from Bioplatforms Australia, enabled by the Commonwealth Government National Collaborative Research Infrastructure Strategy (NCRIS). M.P. and M.A.C are the recipients of the University of Sydney Charles Gilbert Heydon Travelling Fellowship.

## Supporting information captions

**Supplementary Figure S1. Clubroot disease symptoms triggered by Australian P. brassicae isolates in different Brassica host plants.**

*P. brassicae* VIC3 isolate causes clubroot disease symptoms in different Brassica host plants. (B) *P. brassicae* VIC1 and VIC2 isolates cause clubroot disease symptoms in mizuna (*Brassica rapa* var. *niposinica*) plants. In all panels plants were inoculated with spore suspension with a concentration of 1 x 10^7^ spores/ml. Symptom photographs were taken at 35 days post-infection.

**Supplementary Figure S2. Phylogenetic network analysis with a full set of P. brassicae isolates.** A phylogenetic network of all assemblies examined in this study was constructed with SplitsTree and based on a set of 109,960 SNPs produced by the PhAME pipeline. A long branch separates clade A (DAR81272a, DAR8015, DAR44756 and DAR60692) and all other samples in clade B. The tip shapes and colours follow Fig 3, with a network of clade A only presented as Fig 3C.

**Supplementary Table 1.** List of herbaria and field-collected clubroot isolates with their description.

**Supplementary Table 2.** Genome assembly statistics for herbarium isolates sampled in Australia.

