## Supplementary Table 1 for "Phylogenomics and effector analysis of *Plasmodiophora brassicae* genomes unveils a unique and highly divergent clade in Australia"

**Supplementary Table 1.** List of herbaria and field-collected clubroot isolates with their description.

| **Isolate** | **Location** | **Collection date** | **Host** |
| --- | --- | --- | --- |
| VIC1* | VIC, Maffra | 2023-11-28 | *Diplotaxis tenuifolia* |
| VIC2* | VIC, Maffra | 2023-11-28 | *Brassica rapa* var*. niposinica* |
| VIC3* | VIC, Lindenow | 2023-12-12 | *Brassica oleracea* var*. italica* |
| DAR63150 | NSW, Crookwell | 1989-12-11 | *Brassica oleracea* |
| DAR60692 | NSW, Werombi | 1987-09-17 | *Iberis amara* |
| DAR72826 | TAS, Ferry | 1980-04-09 | *Brassica oleracea* |
| DAR44756 | NSW, Glenorie | 1983-06-27 | *Iberis amara* |
| DAR15458 | NSW, Finley | 1965-10-11 | *Sinapis alba* |
| DAR6254 | NSW, Tamworth | 1961-08-18 | *Brassica napus var. napobrassica* |
| DAR6019 | NSW, Penrith | 1960-03-01 | *Brassica oleracea* |
| DAR5008 | NSW, Newcastle | 1958-09-23 | *Lobularia maritima* |
| DAR68422 | SA, Mount Barker | 1992-06-01 | *Brassica rapa subsp. rapa* |
| DAR16316 | NSW, Bathurst | 1967-04-19 | *Brassica oleracea* |
| DAR55183 | NSW, Haymarket | 1985-11-28 | *Brassica rapa var. chinensis* |
| DAR72870 | TAS, Forth | 1980-07-02 | *Brassica rapa* |
| DAR81272a | NSW, Condobolin | 2010-09-10 | *Sinapis sp.* |
| DAR8015 | NSW, Cowra | 2003-09-01 | *Sinapis alba* |

* - isolates from the field collections
