## Supplementary Table 2 for "Phylogenomics and effector analysis of *Plasmodiophora brassicae* genomes unveils a unique and highly divergent clade in Australia"

**Supplementary Table 2.** Genome assembly statistics for herbarium isolates sampled in Australia.

| **Isolate** | **Coverage** | **Assembly size (bp)** | **Number of scaffolds** | **Largest scaffold (bp)** | **N50 length (bp)** | **N90 length (bp)** | **GC content (%)** | **BUSCO (%)** |
| --- | --- | --- | --- | --- | --- | --- | --- | --- |
| DAR63150 | 209 | 23,453,213 | 1314 | 608,646 | 81,899 | 21,481 | 59.57 | 87.45 |
| DAR60692 | 171 | 20,830,876 | 19,494 | 11,982 | 1,167 | 607 | 59.37 | 52 |
| DAR72826 | 167 | 23,299,493 | 945 | 329,326 | 77,873 | 18,282 | 59,57 | 88.24 |
| DAR44756 | 101 | 24,970,396 | 8,059 | 88,734 | 9,647 | 960 | 59.43 | 83.14 |
| DAR15458 | 128* | 23,277,517 | 1,607 | 375,092 | 50,270 | 10,856 | 59.51 | 87.84 |
| DAR6254 | 228* | 24,265,552 | 6,276 | 210,834 | 17,721 | 1,018 | 59.47 | 83.53 |
| DAR6019 | 243 | 23,206,534 | 1,087 | 256,693 | 62,637 | 15,078 | 59.58 | 88.24 |
| DAR5008 | 160* | 23,453,137 | 4,720 | 224,605 | 21,110 | 1,482 | 59.50 | 84.87 |
| DAR68422 | 415 | 23,422,954 | 911 | 454,880 | 80,732 | 21,600 | 59.52 | 88.24 |
| DAR16316 | 279 | 23,208,890 | 1,013 | 256,693 | 69,153 | 16,049 | 59.57 | 88.24 |
| DAR55183 | 310 | 23,211,321 | 1,061 | 256,693 | 64,499 | 15,568 | 59.57 | 88.63 |
| DAR72870 | 255 | 23,498,379 | 1,111 | 365,851 | 63,687 | 15,872 | 59.52 | 87.84 |
| DAR81272a | 279 | 22,859,281 | 3,345 | 246,745 | 24,299 | 2,533 | 59.68 | 83.92 |
| DAR8015 | 279 | 22,781,843 | 4,269 | 102,908 | 17,450 | 1,771 | 59.66 | 85.88 |

* - isolates were sequenced twice to achieve the acceptable coverage. The cumulative coverage of the two runs is provided.
