## Supplementary figures and images for "Phylogenomics and effector analysis of *Plasmodiophora brassicae* genomes unveils a unique and highly divergent clade in Australia"

### Supplementary figure S1

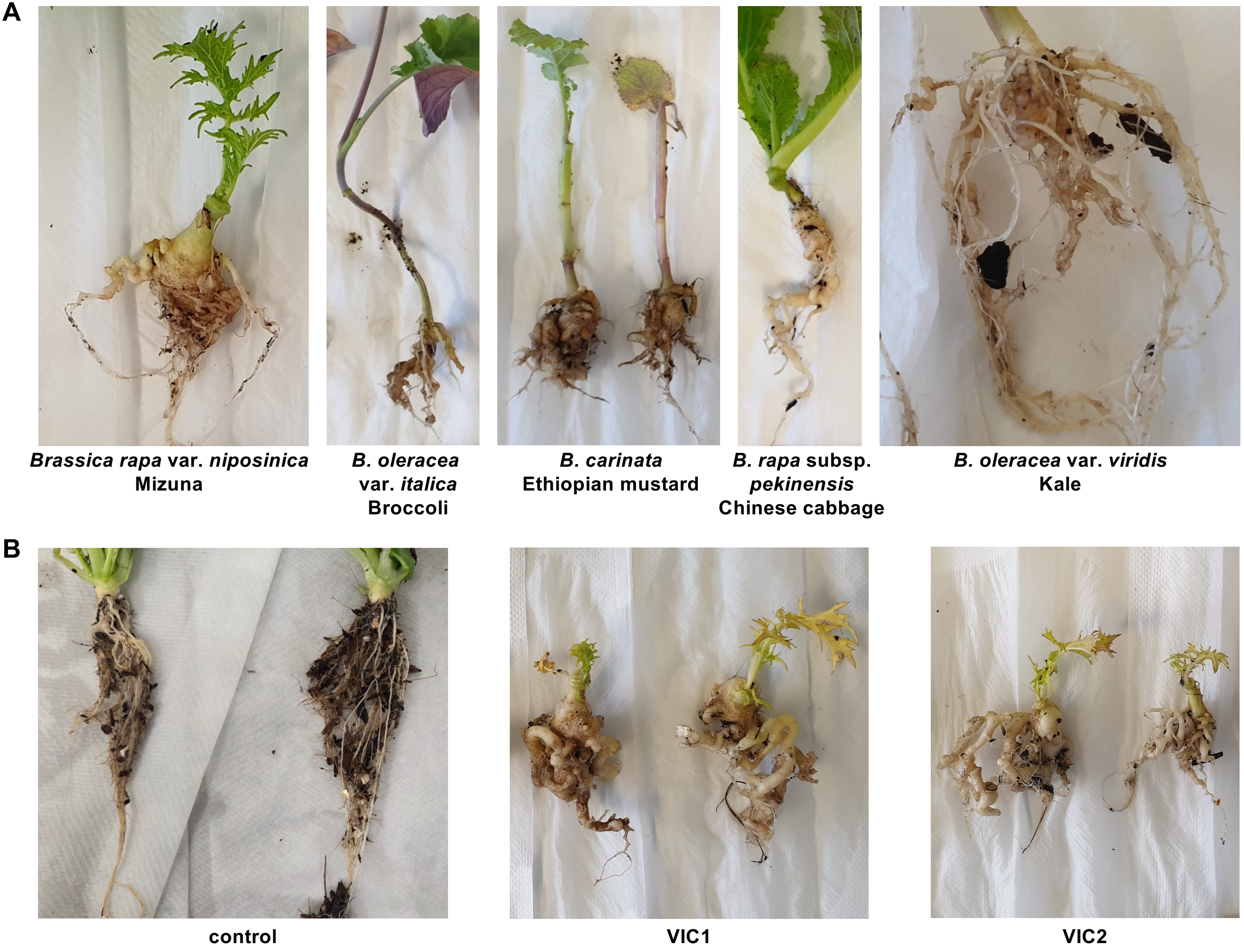

### Supplementary figure S2

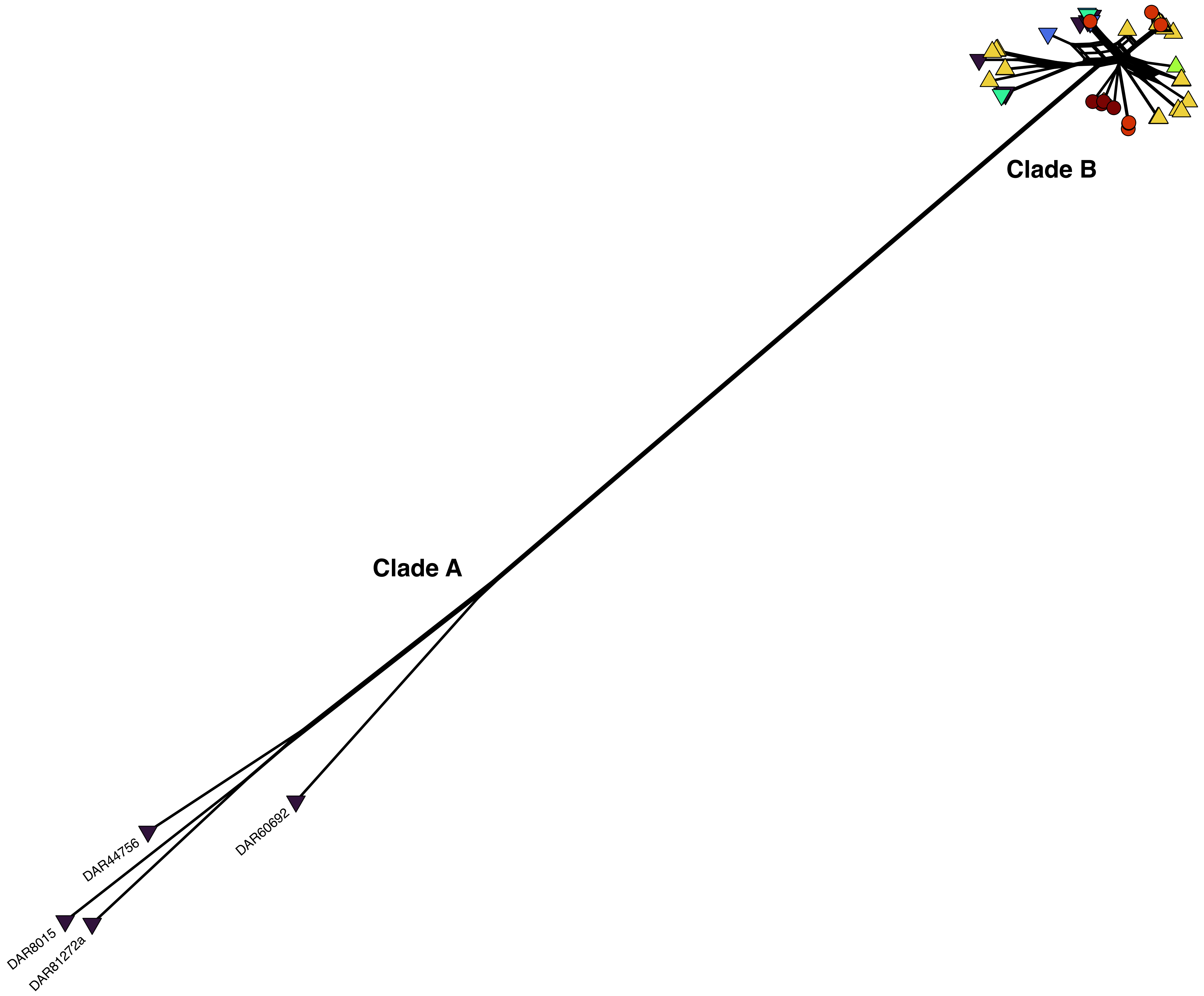
